# A mathematical model of metacarpal subchondral bone adaptation, microdamage, and repair in racehorses

**DOI:** 10.1101/2025.04.06.646486

**Authors:** Michael Pan, Fatemeh Malekipour, Peter Pivonka, Ashleigh V Morrice-West, Jennifer A Flegg, R Chris Whitton, Peta L Hitchens

**Affiliations:** Equine Centre, Melbourne Veterinary School, Faculty of Science, The University of Melbourne, Werribee, VIC 3030, Australia; School of Mathematics and Statistics, The University of Melbourne, Parkville 3010, Victoria, Australia; ARC Centre of Excellence for the Mathematical Analysis of Cellular Systems, University of Melbourne, Parkville, Victoria 3010, Australia; Department of Biomedical Engineering, University of Melbourne, Parkville, VIC 3010, Australia; School of Mechanical, Medical and Process Engineering, Queensland University of Technology, Brisbane, QLD 4000, Australia

## Abstract

Fractures of the distal limb in Thoroughbred racehorses primarily occur because of accumulation of bone microdamage from high-intensity training. Mathematical models of subchondral bone adaptation of the third metacarpal lateral condyles are capable of approximating existing data for Thoroughbred racehorses in training or at rest. To improve upon previous models, we added a dynamic resorption rate and microdamage accumulation and repair processes. Our ordinary differential equation model simulates the coupled processes of bone adaptation and microdamage accumulation, and is calibrated to data on racehorses in training and rest. Sensitivity analyses of our model suggest that joint loads and distances covered per day are among the most significant parameters for predicting microdamage accumulated during training. We also use the model to compare the impact of incremental increasing training programs as horses enter training from a period of rest and maintenance workloads of horses that are race fit on bone adaptation. We find that high-speed training accounts for the majority of damage to the bone. Furthermore, for horses in race training, the estimated rates of bone repair are unable to offset the rate of damage accumulation under a typical Australian racing campaign, highlighting the need for regular rest from training.

**Nomenclature:** 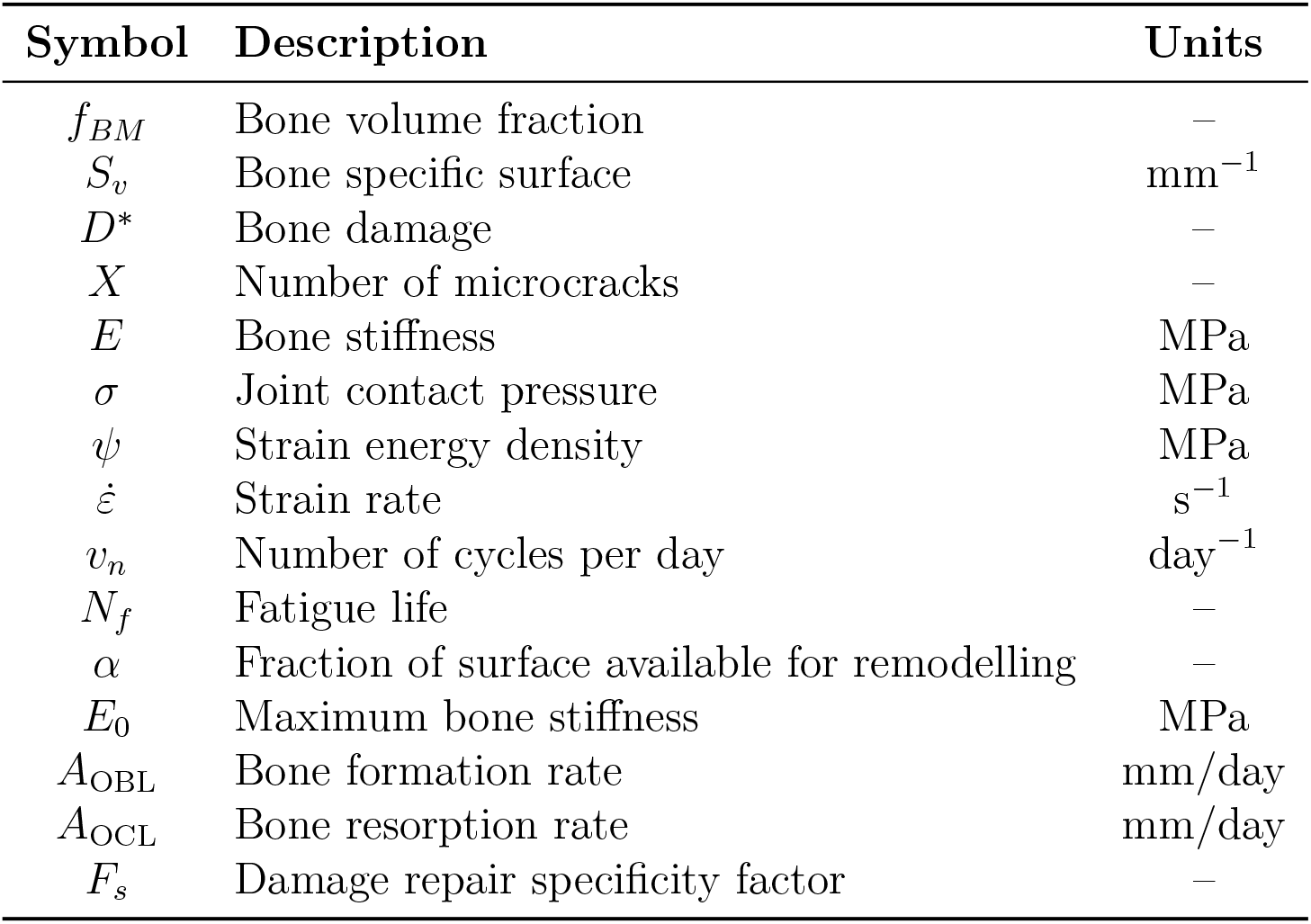

## 1 Introduction

Bone fractures are a major concern in Thoroughbred racehorses, with the subchondral bone of the distal third metacarpal bone being a common site of injury [1–3]. To achieve fitness and musculoskeletal adaptation to race conditions, mechanical loading is primarily introduced via repeated cyclical loading (strides) at incrementally increasing speeds. However, too many cycles at high-intensity loads may be injurious [4–6]. Previous epidemiological studies have found both the speed and distances worked during training to be linked to bone fractures and musculoskeletal injury [6, 7]. However, training can also have a protective effect on bone, with cumulative gallop distances over a career being linked to lower fracture incidence rates [6]. These conflicting findings of both too little and too much high-speed exercise associated with catastrophic musculoskeletal injury [8] highlight the importance of understanding the biological process of bone adaptation, damage accumulation and repair.

Mathematical models of bone adaptation due to mechanical loading have a long history in development and have provided many insights into the fundamental mechanisms playing roles in bone adaptation [9, 10]. In the context of exercise controlled bone adaptation responses in humans two major computational modeling themes have emerged: (i) models for predicting stress fractures in athletes and soldiers [11, 12] and (ii) enhancement of drug efficacy with exercise for osteoporosis treatments [13, 14]. Both of these modeling approaches require linking of the mechanostat feedback with bone damage formation and repair during high cyclic loading and rest intervals, respectively. There have been several modelling studies on bone adaptation and damage in the trabecular and cortical bone of humans [12, 15–17].

While the above-described models of coupled bone adaptation, damage formation and repair have been applied to human studies, to the authors’ knowledge, no such models exist for Thoroughbred racehorses. A previous model of subchondral bone adaptation in the third metacarpal bone in Thoroughbred racehorses predicted changes in bone volume fraction (*f*_*BM*_) with joint stress [18]. Morrice-West et al. [19] subsequently developed a method for calculating bone damage accumulated using experimentally derived relationships between stress and bone fatigue life [20].

Here, we aim to extend our previous model of subchondral bone adaptation in the third metacarpal bone in Thoroughbred racehorses [18] to incorporate bone damage formation and repair processes. In particular, we add a dynamic resorption rate which depends on loading intensity and differential equations accounting for bone microdamage formation and repair. The resorption rate of the third metacarpal (MCIII) condyles is generally higher than the bone formation rate except at the highest levels of mechanical loading [21–23]. Hitchens et al. [18] modelled bone resorption as a constant rate, however, osteoclastic resorption is inhibited by intense race training [24, 25]. The rate of bone damage accumulation increases exponentially with the magnitude of load [19, 20], and we use this relationship to model damage formation in our model. Furthermore, we derive equations for bone repair that are coupled to the dynamics of bone formation and resorption following the approach of Martin [15] and Hazelwood et al. [26].

In this study, we aim to develop a computational model of bone adaptation for Thoroughbred racehorses, including damage formation and repair. We calibrate the model to measurements from horses resting and in race training, and simulate the response of bone to a typical training program based on surveyed workloads of racehorses in Victoria, Australia [27]. We hypothesise that (i) the joint stress and number of cycles (strides) per day are key determinants of bone injury risk; (ii) training programs with greater volumes of high-speed gallops are more likely to lead to bone injuries; and (iii) rest periods are essential to bone repair. This extended model will help us to enhance understanding of the processes of bone adaptation, microdamage formation, and repair to the subchondral bone of the distal metacarpus of racehorses, forming the basis for assessing and developing safer training strategies that reduce the risk of racehorse injury.

## 2 Methods

In this section, we describe our model of bone adaptation and damage within a region of interest at the lateral condyle of the third metacarpal bone (Figure 1), which experiences high load and is a common site of subchondral bone damage [24, 29]. The model is defined by Eqs (1)–(12). A summary of the processes modelled is schematically shown in Figure 2.

**Figure 1:**
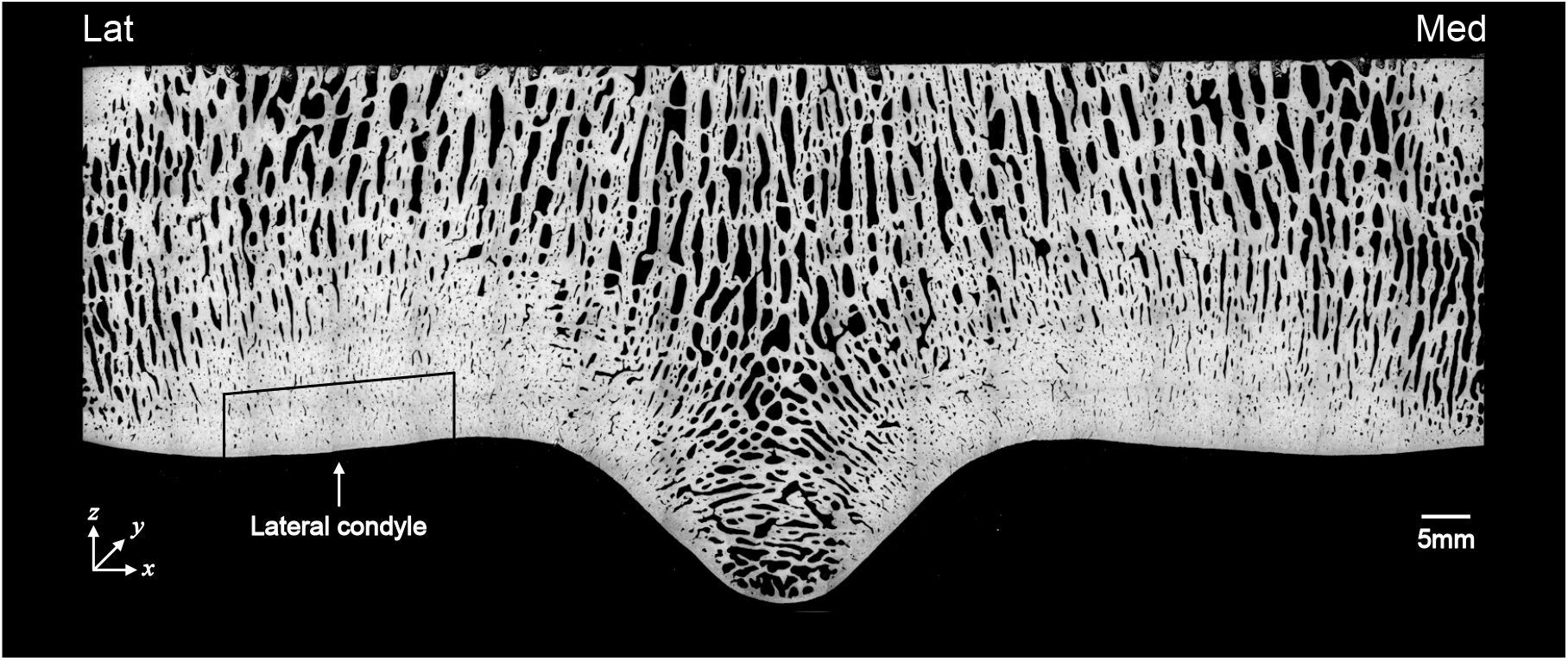
Back-scattered electron microscopy image of the distal third metacarpal bone in a Thorough-bred racehorse. The volume of interest is the mid lateral condyle articular margin, indicated by the arrow. The coordinate system is defined as *x*, mediolateral; *y*, dorsodistal *z*, distopalmar direction. Adapted with permission from Holmes et al. [24].

**Figure 2:**
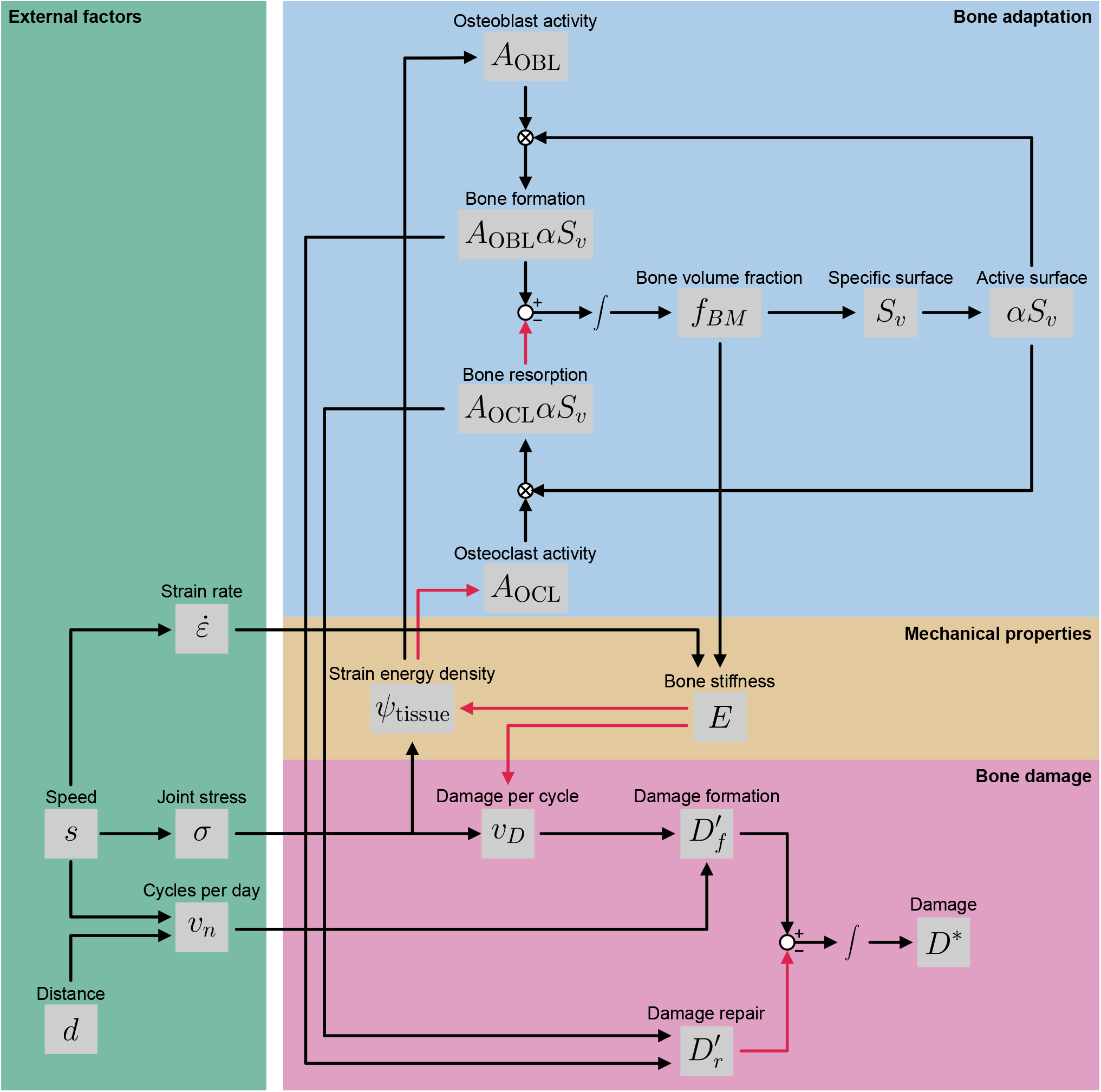
Schematic of the bone adaptation and damage model. Activating relationships are shown by black arrows, and inhibitory relationships are shown by red arrows. (Blue box) The subchondral bone consists of regions of bone matrix and empty regions of marrow. Remodelling occurs at the interface of bone and marrow. On average, the surface area (*S*_*v*_) available for remodelling decreases with increasing bone volume fraction (*f*_BM_). Osteoblasts (with activity *A*_OBL_) are responsible for bone formation, whereas osteoclasts (with activity *A*_OCL_) are responsible for bone resorption. (Yellow box) The bone senses mechanical stresses through its strain energy density (*ψ*_tissue_), which acts as a signal for bone adaptation. (Pink box) Stresses incurred on the bone lead to the accumulation of damage (*D*^*∗*^) through microcracks. These microcracks can be removed through osteoclast resorption. (Green box) The speeds (*s*) and distances (*d*) of training programs correspond to both joint stress (*σ*) and number of cycles (*v*_*n*_) [28], inducing bone formation but also causing microdamage. Open circles indicate the addition or subtraction of variables, and circles with crosses inside indicate multiplication.

### 2.1 Base mathematical model of bone adaptation

The base model of subchondral bone adaptation of the equine distal metacarpus is described in detail in Hitchens et al. [18]. The model consists of an ordinary differential equation (ODE) described by two main equations

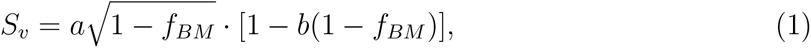

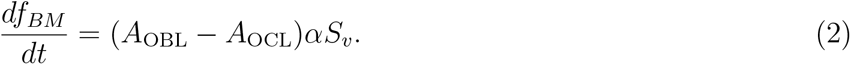

Eq. (1) is a function that expresses the relationship between the bone volume fraction *f*_*BM*_ [dimensionless] and bone specific surface *S*_*v*_ [mm^*−*1^] (Figure 2; blue box). This equation is the square root function previously derived by Lerebours et al. [30], with parameters modified to fit bone data from racehorses (*a* = 11.42 mm^*−*1^ and *b* = −0.02). Eq. (2) is a mass balance of the bone volume fraction [31] (Figure 2; blue box). *A*_OBL_ [mm/day] is the bone formation activity (bone volume change per unit time per unit surface area), and *A*_OCL_ [mm/day] is the bone resorption activity. In Hitchens et al. [18], *A*_OBL_ is dynamic and dependent on strain energy density *ψ*_tissue_ [MPa], whereas *A*_OCL_ was assumed constant. However, as explained later in Subsection 2.2, we use dynamic rates for both activities, and update the equations for both *A*_OBL_ and *A*_OCL_. The rates of bone formation and resorption are both proportional to *α*, the specific surface available for bone remodelling. It is assumed that bone properties are homogeneous throughout the metacarpal condyles.

The model also assumes that osteocytes sense mechanical forces through the strain energy density *ψ*_tissue_, which in turn affects the osteoblast activity *A*_OBL_ (Figure 2; yellow box). Supposing there is a uniaxial linear constitutive relationship between the joint stress *σ* [MPa] and strain *ε*, the strain energy density is given by

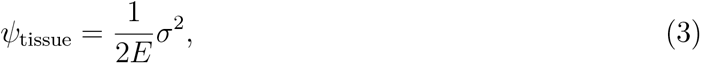

where *E* [MPa] is the tissue stiffness [18]. The loading state is assumed to be uniaxial, i.e. *σ* = *σ*_*zz*_ and *σ*_*xx*_ = *σ*_*yy*_ = *σ*_*xy*_ = *σ*_*xz*_ = *σ*_*yz*_ = 0 (coordinates indicated in Figure 1). Since bone stiffness has been observed to increase with both bone volume fraction and strain rate, the tissue stiffness is set to

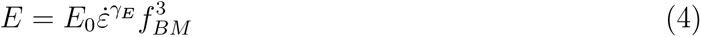

with *γ*_*E*_ = 0.06, as described previously [18].

In addition to the joint stress, the number of cycles (strides) per day can also influence bone adaptation [32]. However, we have not modelled this relationship as bone formation has been observed to plateau above 36 cycles per day [32], and we assume that horses train at distances above this threshold, given that horses galloping at a typical volume of 1800 m per week will accumulate approximately 40 cycles per day.

### 2.2 Extended mathematical model of bone adaptation

The base model is updated to incorporate the effects of *ψ*_tissue_ on both the bone formation activity *A*_OBL_ and resorption activity *A*_OCL_ (Figure 2; blue box). Osteocytes in the bone tissue are mechanosensors, that is, they are assumed to sense a mechanical load in the form of strain energy density. These osteocytes release biochemical signals that affect bone formation and resorption rates. Bone formation rates are higher in unadapted bone subjected to high-speed exercise [21, 23]. This is modelled using the Hill equation

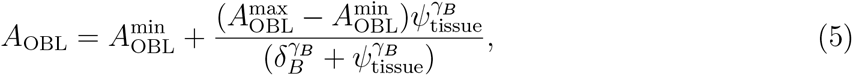

where 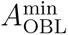 [mm/day] is the minimum bone formation rate, 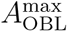 [mm/day] is the maximum bone formation rate, *δ*_*B*_ [MPa] is the half-saturation constant and *γ*_*B*_ [dimensionless] is the sigmoidicity.

Conversely, high-speed exercise inhibits bone resorption [24, 33, 34]. Thus, the bone resorption rate decreases when bone is subjected to higher loads. This is modelled using the decreasing Hill function

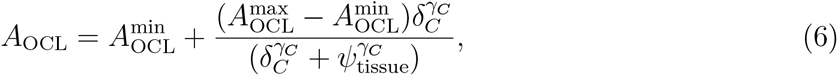

with constants defined similarly to Eq. (5). We set the minimum resorption rate to 0.001 mm/day [35], and assume that *γ*_*C*_ = 2 to achieve a sigmoidal relationship. Horses at rest have been observed to have four times the erosion surface of horses in full training, which translates to higher resorption rates [24]. We thus tune this equation by selecting a proxy for rest at a load of 30 MPa, and for race-fit training and racing at 90 MPa. These values were taken from Hitchens et al. [18], and are consistent with estimated joint loads for walking and trotting [36] as well as loads applied during mechanical tests on subchondral bone [20], based on the yield stresses measured in lateral condyles [37]. Using the corresponding bone stiffnesses for a horse that has been adapted to racing speeds and distances at *f*_*BM*_ = 0.9 (see Table 3 of Hitchens et al. [18]), these stresses correspond to strain energy densities of 0.31 and 2.36 respectively. We therefore set the half-saturation constant to *δ*_*C*_ = 1.0 MPa so that bone resorption is activated in between the two loads. While the maximum resorption rate 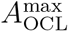 is estimated to be as high as 0.011 mm/day in untrained two-year-old horses [35], we believe this is an overestimate as it results in excessive bone resorption within our model. Thus, we fit this parameter to time-series data on bone volume fractions (Subsection A.3, electronic supplementary material). The curves for *A*_OBL_ and *A*_OCL_ (after fitting) as functions of *ψ*_tissue_ are shown in Figure S8 of the electronic supplementary material.

### 2.3 Bone microdamage

It is widely accepted that the majority of equine bone injuries occur due to bone fatigue, the gradual accumulation of microcracks in the bone until a fracture occurs. Microdamage formation as a result of cyclical loading is non-linear, typically occurring in three stages. In a study of racehorses, most subchondral bone specimens under compressive fatigue increased stiffness rapidly in the initial stage, then plateaued during the mid-stages close to or at maximum stiffness, then in the final stage decreased rapidly to failure [20]. Less commonly, in specimens with shorter fatigue life, either stiffness decreased sharply between the initial stage and the plateau phase or a steady decrease in stiffness replaced the plateau stage [20]. The reason for the plateau in stiffness and strength is due to build up of residual strain. Eventually, the microcracks propagate and overwhelm the bone repair process until catastrophic failure [38]. The loads experienced by the subchondral bone of the distal third metacarpal bone within Thoroughbred racehorses are predominantly compressive [39], leading to microcracks oriented obliquely to the articular surface [40].

Carter and Caler [41] used the framework of continuum damage mechanics to develop a mathematical model which linked load and bone damage. Subsequently, several researchers expanded upon this model [12, 15, 42]. While there are multiple definitions of damage in the literature [13, 15, 43, 43–46], we define damage *D*^*∗*^ as the life fraction *D*^*∗*^ = *n/N*_*f*_, where *n* is the number of cycles at a given stress and *N*_*f*_ is the fatigue life, the number of cycles a volume of bone can resist prior to failure. Thus, the rate of damage accumulation is

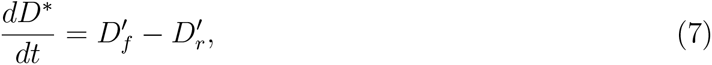

where 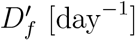 is the rate of damage formation and 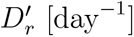 is the rate of damage repair (Figure 2; pink box).

From the definition of the damage *D*^*∗*^, the bone is considered to have failed at *D*^*∗*^ = 1, which corresponds to the failure of pocket of bone within the lateral condyle of the third metacarpal bone through excessive accumulation of microdamage. We note that this in most cases does not lead to a gross fracture of bone, but will arise as palmar osteochondral disease, which is commonly observed in active racehorses [29].

#### 2.3.1 Damage formation

Damage is incurred with each loading cycle (stride) and accumulates at faster rates at greater loads. The rate of damage formation 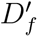 can therefore be expressed using the equation

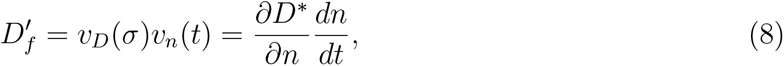

*D*where *v*_*D*_(*σ*) = *∂D*^*∗*^*/∂n* is the damage incurred per cycle and *v*_*n*_(*t*) = *dn/dt* is the number of cycles per day [12].

The ability of bone to resist fatigue at a particular stress level is characterised by the fatigue life *N*_*f*_. Higher stresses impart greater damage to the bone, hypothesised to be due to the increased dissipation of mechanical energy into the bone [47]. Mechanical testing on equine subchondral bone has previously shown that the fatigue life of bone is related to the compressive stress (i.e. positive *σ* corresponds to compression) of cycling through a power law:

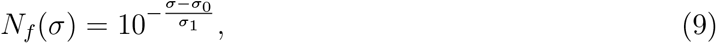

where *σ*_0_ = 134.2 MPa and *σ*_1_ = 14.1 MPa [20]. Bone stiffness provides resistance to bone fatigue, thus, we adjust the fatigue life to include an additional multiplicative factor dependent on the Young’s modulus [15, 43]:

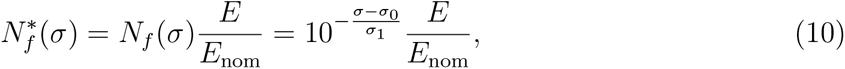

where *E*_nom_ is a reference stiffness. Given that Eq. (9) was derived from racehorses in training, we assume that *E*_nom_ = *E*_0_·0.9^3^·0.36^0.06^ = 1714.1 MPa, i.e. the experiments were conducted on bone with *f*_*BM*_ = 0.9 and a strain rate of 0.36 s^*−*1^.

Since the bone fails at *D*^*∗*^ = 1, the damage incurred per cycle is

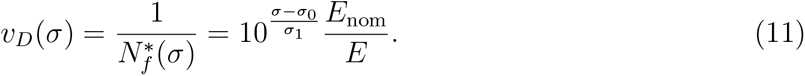

While bone stiffness transiently increases before gradually decreasing over its fatigue lifetime [20], we have not included this phenomenon in the model for simplicity.

#### 2.3.2 Damage repair

Following the incursion of microdamage to bone, microcracks are removed by the resorption of bone by osteoclasts and subsequent bone formation (Figure 2; pink box). Bone remodelling occurs within bone multicellular units (BMUs) that contain populations of both osteoblasts and osteoclasts. Though microdamage is more likely to occur focally at loading sites [48], we model microdamage as an even distribution throughout the entire volume of subchondral bone. Thus, the damage repair term in Eq. (7) is

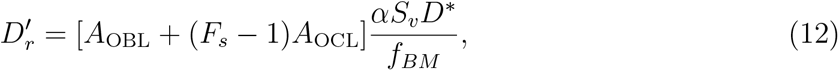

with a full derivation provided in Subsection A.1 of the electronic supplementary material. Briefly, bone formation (*A*_OBL_) leads to a reduction of damage due to the dilution of microcracks across a larger volume. Bone resorption also leads to damage repair if osteoclasts target microcracks specifically. It has been observed that bone resorption is more likely to occur at sites of microcracks [49, 50]. Accordingly, we include a damage repair specificity factor, *F*_*s*_ [dimensionless], which is the rate at which bone resorption sites co-localise with microcracks relative to the random emergence of resorption sites [15]. Thus, *F*_*s*_ = 1 corresponds to untargeted resorption. Following Martin [15], we set *F*_*s*_ = 5, since cracks are estimated to be 4–6 times more likely to be associated with resorption sites compared to chance alone [49, 50]. It is worth noting that while *F*_*s*_ increases the extent of bone repair, it does not increase the rate of bone resorption.

## 3 Results

### 3.1 Comparison of mathematical model to data

The model is calibrated to cross-sectional data on bone volume fraction and the median time in training for horses that sustained fractures [18, 24, 51, 52] (details in Subsection A.3, electronic supplementary material). Parameter estimation is performed using optimisation [53, 54], and comparisons of model simulations to data are shown in Figure 3. The response of bone to training and rest are depicted in Figures 3a and 3b respectively. Despite the marked variance within the data, the model captures trends including the gradual increase in bone volume fraction with training and more rapid loss of bone in response to rest. Furthermore, the model faithfully fits the median time to fracture (133 days) under a high workload (Figure 3c). The parameters used in the model (including fitted parameters) are given in Table 1.

**Table 1:** Parameters in the mathematical model of subchondral bone adaptation in Thoroughbred racehorses, and their ranges for sensitivity analysis.

| Parameter | Description | Units | Value [Initial (fitted)] | Range | Reference |
| --- | --- | --- | --- | --- | --- |
| $\sigma$ | Joint contact pressure | MPa | 81.1 | [30.0, 102.0] | <sup>a</sup> |
| $\dot{\epsilon}$ | Strain rate | s<sup>-1</sup> | 0.2974 | [0.05, 0.46] | <sup>a</sup> |
| $v_n$ | Number of cycles per day | day<sup>-1</sup> | 43.7 | [25.0, 120.0] | <sup>a</sup> |
| $\alpha$ | Fraction of specific surface | – | 0.19 | [0.095, 0.285] | Whitton et al. [48] |
| $a$ | Fitting parameter relating $f_{BM}$ to $S_v$ | mm<sup>-1</sup> | 11.42 | [5.71, 17.13] | Hitchens et al. [18] |
| $b$ | Fitting parameter relating $f_{BM}$ to $S_v$ | – | −0.02 | [−0.2, 0.2] | Hitchens et al. [18] |
| $E_0$ | Maximum bone stiffness | MPa | 2500 | [1250.0, 3750.0] | Martig et al. [20] |
| $\gamma_E$ | Exponent of strain rate in stiffness equation | – | 0.06 | [0.03, 0.09] | Carter and Hayes [55] |
| $A_{OBL}^{\min}$ | Minimum bone formation rate | mm/day | 0.001094 | [0.0, 0.00301] | Firth et al. [22] |
| $A_{OBL}^{\max}$ | Maximum bone formation rate | mm/day | 0.0127 (0.00603) | [0.00301, 0.00904] | Davies [21] |
| $\delta_B$ | Half-saturation stress of bone formation | MPa | 7 | [3.5, 10.5] | Les et al. [56] |
| $\gamma_B$ | Sigmoidicity of bone formation | – | 1 | [1.0, 5.0] | Hitchens et al. [18] |
| $A_{OCL}^{\min}$ | Minimum bone resorption rate | mm/day | 0.001 | [0.0, 0.0018] | Boyde and Firth [35] |
| $A_{OCL}^{\max}$ | Maximum bone resorption rate | mm/day | 0.011 (0.00358) | [0.0018, 0.0054] | Boyde and Firth [35] |
| $\delta_C$ | Half-saturation stress of bone resorption | MPa | 1 | [0.5, 1.5] | <sup>b</sup> |
| $\gamma_C$ | Sigmoidicity of bone resorption | – | 2 | [1.0, 5.0] | <sup>b</sup> |
| $\sigma_0$ | Fitting parameter relating stress to fatigue life | MPa | 134.2 (139.0) | [125.1, 152.9] | Martig et al. [20] |
| $\sigma_1$ | Fitting parameter relating stress to fatigue life | MPa | 14.1 | [12.69, 15.51] | Martig et al. [20] |
| $E_{\text{nom}}$ | Reference stiffness for stress-fatigue life equation | MPa | 1714.1 | [857.0, 2571.2] | <sup>c</sup> |
| $F_s$ | Damage repair specificity factor | – | 5 | [1.0, 10.0] | Martin [15] |

<sup>a</sup> Calculated from equations in Subsection A.2 of the electronic supplementary material with a speed of 14.0 m/s and a distance of 8000m every 30 days (Table 2, programme group 2 of Morrice-West et al. [27]). <sup>b</sup> Selected to achieve four times higher resorption in full training relative to rest [24]. <sup>c</sup> Calculated assuming $f_{BM} = 0.9$ .

**Figure 3:**
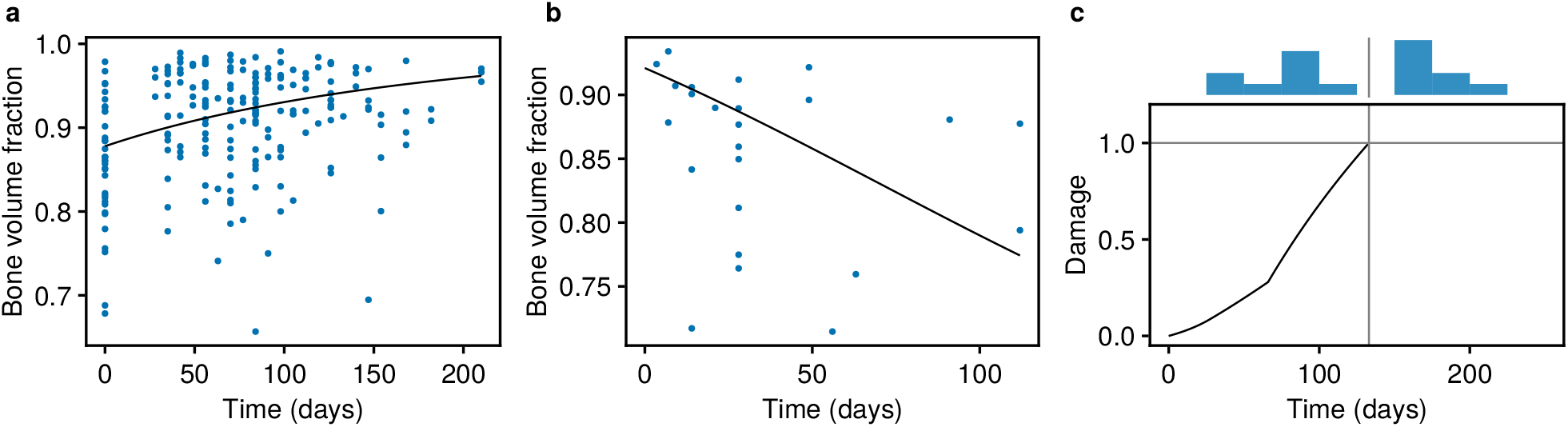
Comparisons of model output to bone volume fraction and time to fracture data [18]. The mathematical model has been fit to data (details in Subsection A.3, electronic supplementary material), with parameter values in Table 1. (a) Adaptation of bone volume fraction (*f*_*BM*_) in response to training, compared to measurements from *n* = 213 limbs from 87 horses; (b) De-adaptation of bone volume fraction to rest, compared to *n* = 24 limbs from 24 horses. For (a) and (b), black lines indicate model simulations, and blue dots indicate data points. (c) Simulation of a fracture at high workloads, where the fracture occurs when the damage *D*^*∗*^ reaches 1, at which point the simulation is terminated. A histogram of the duration of preparations prior to fracture is shown above the main plot, with the median indicated with a vertical grey line. Data were collected from *n* = 16 horses.

### 3.2 Sensitivity of bone volume fraction and damage to parameters

Initial simulations of the calibrated model under varying stresses *σ* indicate that both bone volume fraction and damage increase in response to higher joint stresses (Subsection B.1, electronic supplementary material). To generalise this analysis to all model parameters, we use a sensitivity analysis (details in Subsection A.4, electronic supplementary material) to identify the parameters that produce the greatest changes in bone volume fraction and damage. Figure 4 shows the results from this sensitivity analysis, using the partial rank correlation coefficient (PRCC) as the sensitivity measure [57]. Under this measure, parameters with very positive (negative) sensitivities tend to increase (decrease) the output variable most consistently. The parameters *σ*, 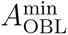 and 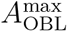 are among those that increase *f*_*BM*_ the most consistently (Figure 4a). This is expected, due to the role of stress and osteoblast activity in bone formation. The surface area parameters *a* and *α* also have a tendency to increase bone volume fraction, although this may be because our simulations are more likely to correspond to bone formation than resorption. The bone stiffness *E*_0_ is the parameter most consistently associated with loss of bone, followed by 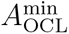, *δ*_*B*_ and *γ*_*B*_. The bone stiffness *E*_0_ has a negative PRCC because it inhibits the mechanical signal required for bone formation (Eq. (3)). Unsurprisingly, increasing the rate of bone resorption through 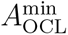 leads to a loss of bone mass. The parameters *δ*_*B*_ and *γ*_*B*_ appear to reduce the bone formation rates at low strain energy densities, and as a result lead to a net loss of bone mass.

**Figure 4:**
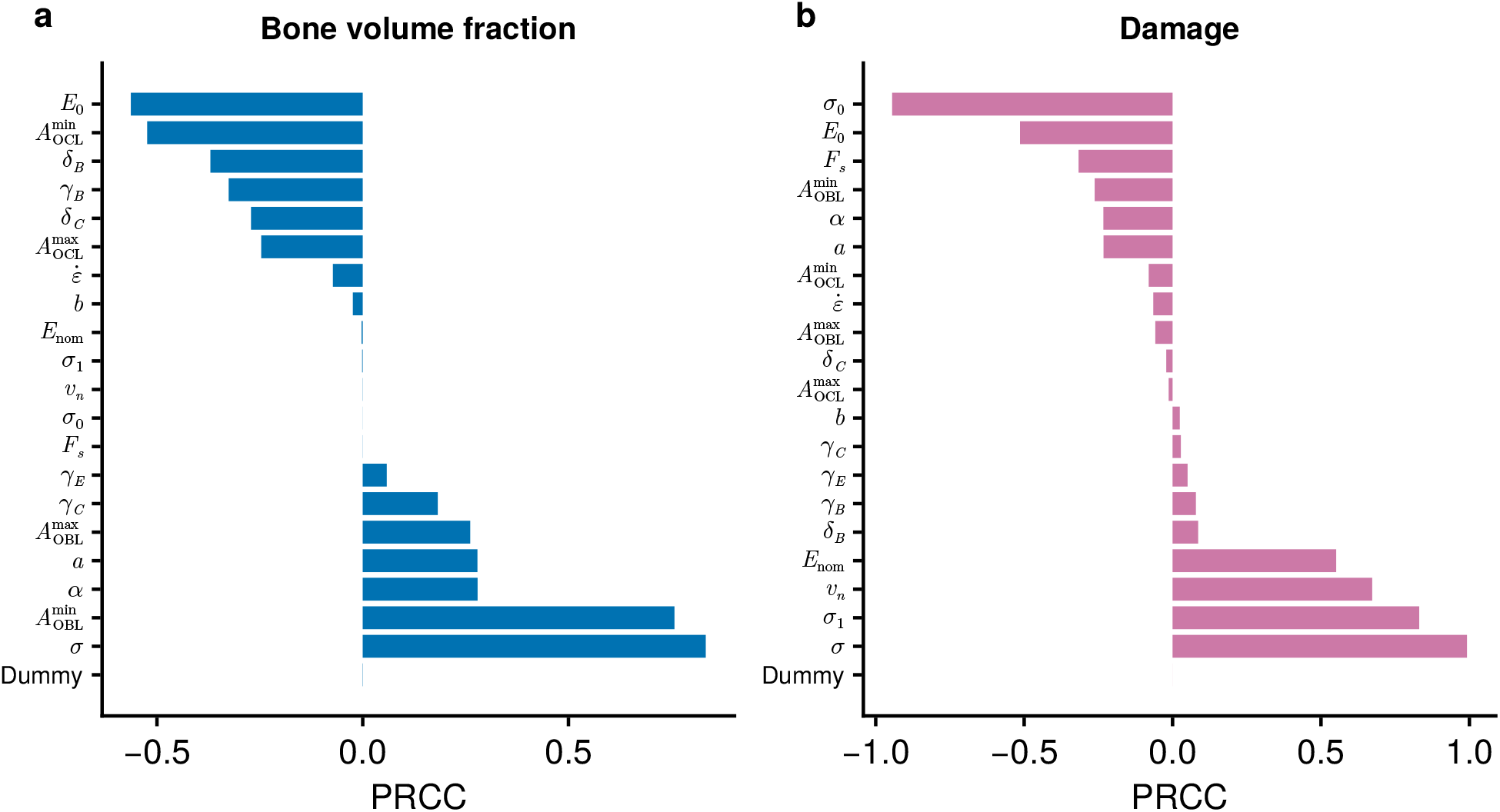
Sensitivity analysis of parameters using the partial rank correlation coefficient (PRCC). Effects of each parameter on (a) bone volume fraction *f*_*BM*_; and (b) damage *D*^*∗*^ after 10 weeks. PRCC values of high magnitude indicate parameters with a consistent influence on the output variables of interest, with the sign indicating the direction of correlation. The model described by Eqs (1)–(12) is simulated using an initial condition of *f*_*BM*_ = 0.7 and *D*^*∗*^ = 0. Descriptions of parameters and their ranges are shown in Table 1. A dummy variable with no effect on the output has been included to detect potentially spurious relationships.

The stress *σ* and cycles per day *v*_*n*_ are among the most consistent inducers of bone damage *D*^*∗*^ (Figure 4b). Other parameters that tend to increase bone damage are *σ*_1_ and *E*_nom_. The parameter most associated with reduced damage is *σ*_0_, due to its ability to offset stresses in the exponential relationship between stress and fatigue life (Eq. (9)). *E*_0_ and *F*_*s*_ are also able to reduce bone damage, indicating the role that bone stiffness and microdamage resorption play in resisting failure.

While the PRCC reliably estimates the manner in which a parameter affects an output, it may fail to capture the relative magnitude of these effects [58]. The Sobol index is an alternative parameter sensitivity measure that accounts for the magnitude of changes [59, 60]. In line with the PRCC, results for Sobol indices show that the stress is a major factor contributing to bone volume fraction and damage, and that cycles per day has a strong influence on damage (Subsection B.2, electronic supplementary material). The sensitivities for most parameters are largely independent of time (Figures S9–S10, electronic supplementary material).

### 3.3 Temporal response of bone to training programs

The sensitivity analysis only considers training programs that are constant in time. To account for changes in training regimes as racehorses progress from rest to racing, we model the response of bone to a typical Australian training and racing preparation [27, 61–64], as summarised in Figure 5 (details in Subsection A.2, electronic supplementary material). We assume horses start training in their 2-year-old season (in the southern hemisphere, after 1 August in the year that they turn 2-years-old) with an initial bone volume fraction of *f*_*BM*_ = 0.61 and zero damage. While two-year-old horses are trained at smaller distances compared to mature horses [27], we have assumed for simplicity that training programs are constant over a horse’s career. The bone is simulated over four preparations consisting of rest, pre-training, progressive training and race-fit training, and results are shown in Figure 6. Over the first preparation, the bone volume fraction increases to 0.85. While horses are exposed to higher loads during the race-fit period, the higher bone volume fraction limits the surface area available for remodelling, and ultimately leads to lower bone formation compared to the preceding progressive period. The bone volume fraction gradually increases from preparation to preparation, but varies between 0.86 and 0.93 during the fourth preparation, consistent with values seen in horses at rest and training respectively [20]. The damage corresponding to this training program rises during both progressive training and race-fit training, to a value of *D*^*∗*^ = 0.90 by the end of the first racing preparation. The damage is repaired during the subsequent rest period, although the repair processes are unable to completely remove this damage. While progressive training resumes before the damage has completely repaired, the maximum damage attained in subsequent preparations is approximately equal to that at the end of the first preparation.

**Figure 5:**
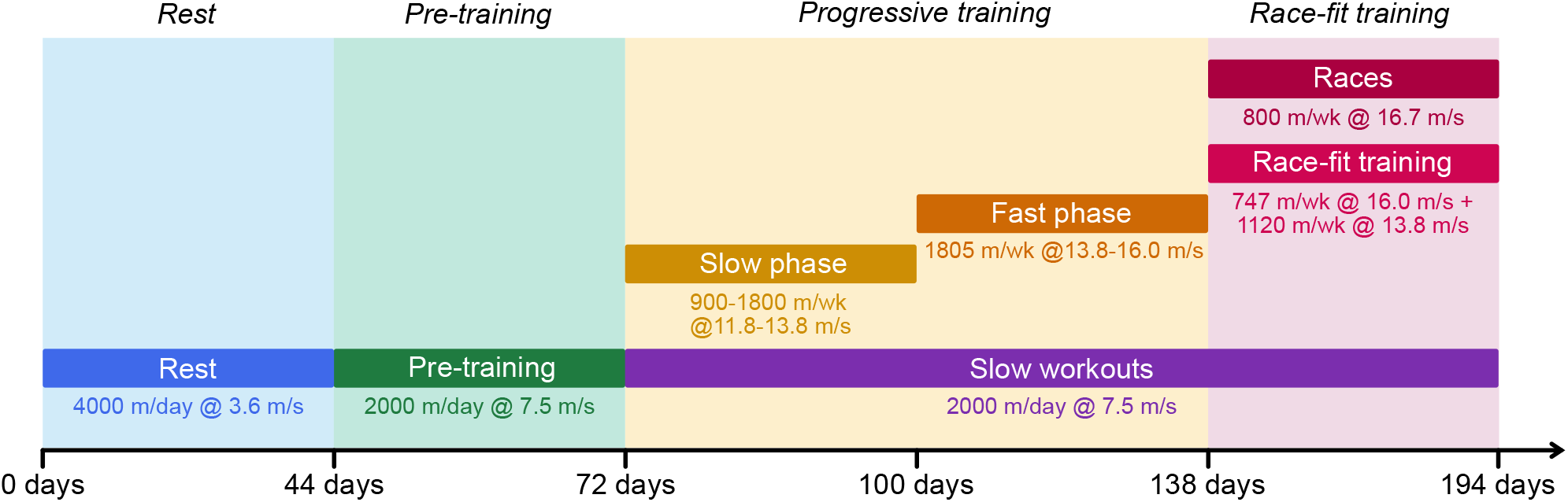
Schematic of the simulated training program (*‘preparation’*). Note that time starts from the start of a rest period and is not shown to scale. Distances for longer-distance training at canter speeds and below are expressed in distances per day, whereas faster workouts (which are not performed every day) are expressed in distances per week.

**Figure 6:**
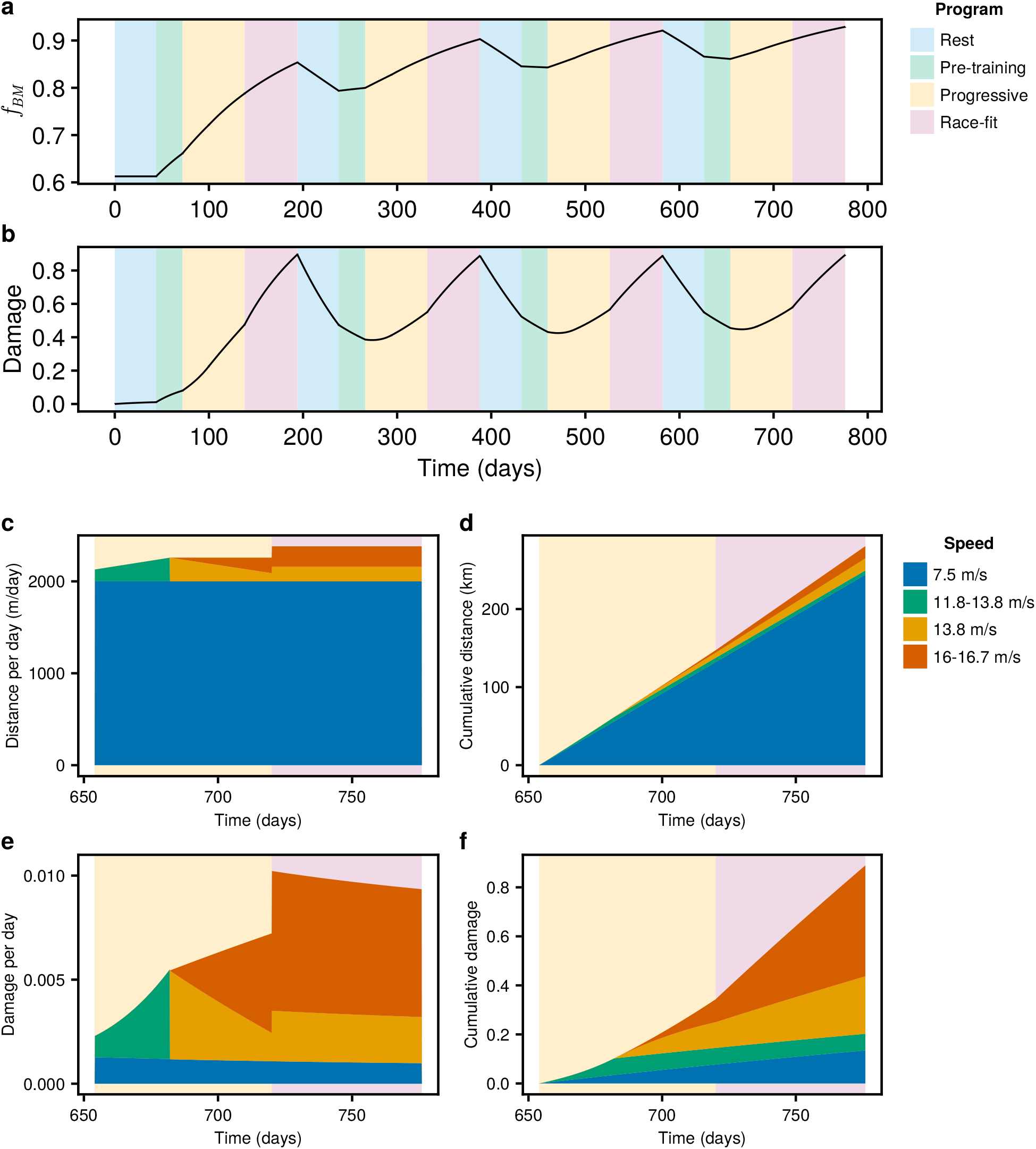
Dynamic simulations of bone in response to a typical training program in Victoria, Australia, as defined in Figure 5. The background colours indicate periods of rest (blue), pre-training (green), progressive training (yellow) and race-fit training (pink). The training program is described in Subsection A.2 of the electronic supplementary material, with speeds and distances covered during training defined in Subsection A.2.2. Simulations are run starting from *f*_*BM*_ = 0.61 and *D*^*∗*^ = 0, corresponding to an untrained racehorse. The simulation is run over four preparations. Changes in (a) bone volume fraction *f*_*BM*_ and (b) damage *D*^*∗*^ are plotted against time. During the final (fourth) instance of progressive and race-fit training (*t∈* [654, 776]), the distance and damage incurred at speeds *≥* 7.5 m*/*s are summarised in (c–f). (c) Distance per day; (d) cumulative distance; (e) damage per day; and (f) cumulative damage.

A breakdown of the distances accumulated during the final (fourth) progressive and race-fit training preparations is plotted in Figure 6c. The distances are coloured according to the corresponding speeds. There is a gradual increase in distances covered at the highest speed of 16.0 m/s as the horse shifts from progressive training (yellow background) to racing (pink background). By the end of the racing period, the horse has covered 280.3 km, with 243.8 km at canter (7.5 m/s), 15.6km at racing speed (16.0–16.7 m/s), 15.5 km at a fast gallop (13.8 m/s) and 5.4 km at slow gallop (11.8–13.8 m/s) (Figure 6d). The damage incurred per day is plotted in Figure 6e, and the resulting cumulative damage is shown in Figure 6f. High speeds above 13.8 m/s account for the majority of damage incurred during training (Figure 6f).

## 4 Discussion

In this study, we present a new mathematical model coupling the dynamics of bone adaptation and bone damage in the subchondral bone of the distal third metacarpus of Thoroughbred racehorses. Our model expands on a previous model of bone adaptation [18] through the addition of a dynamic resorption rate (in addition to a dynamic deposition rate) and variables accounting for bone damage. The coupling of bone adaptation and damage dynamics enables new analyses of the response of bone to training. This includes a sensitivity analysis of the parameters most relevant for bone damage, and estimates of how bone damage dynamically evolves in response to various phases of training.

Our model is consistent with data relating to both bone volume fraction and median times to fracture during a racing preparation (Figure 3). Our previous model was already consistent with bone volume fraction [18], and our fits in this current work confirm that bone resorption at rest occurs at faster rates than bone formation in training (in our model, *f*_*BM*_ increases by 0.053 in response to 100 days of training in Figure 3a, but decreases by 0.131 in response to 100 days of rest in Figure 3b). However, the two models differ in the time taken for a steady state to be achieved during rest (over 16 weeks, compared to 4–10 weeks in Hitchens et al. [18]) (Figure 3b). Further investigation is required to better characterise de-adaptation times in racehorses. In addition to fitting changes in bone volume fraction, we fit our model to data on horses that had sustained a fracture [18], observing a time to failure consistent with their time in training. However, we note that this fit requires revising our estimate of *σ*_0_ from 134.2 MPa to 139.0 MPa (Table 1). From Eq. (11), this corresponds to an estimated 54.5% reduction in damage formation rate compared to the relationship derived by Martig et al. [20]. We speculate that this discrepancy is due to differences in forces that bones experience *in vitro* compared to *in vivo*. Martig et al. [20] performed unconfined mechanical tests on subchondral bone cylinders extracted from the lateral condyle of the third metacarpal bone. However, the region of interest in reality is surrounded by bone, connective tissue and overlying cartilage *in vivo* [65]. Thus, given the same externally applied load, we expect that these surrounding structures would dissipate some of the force, leading to lower stresses experienced by the subchondral bone region of interest and mitigating the extent of damage incurred. The application of finite-element models of the metacarpophalangeal joint may provide a means for accounting for these effects [36], but has yet to be applied to galloping horses.

Our sensitivity analysis identifies key parameters affecting bone volume fraction and induction of bone damage. We find that the stress *σ* is a major factor contributing to both bone formation and damage, with positive PRCC values for both outputs (Figure 4). These results are consistent with previous studies on racehorses, which find that (i) training is associated with higher bone volume fractions in racehorses [24, 35]; (ii) high acute workloads are associated with higher rates of musculoskeletal injuries [6]; and (iii) bone failure is accelerated by the application of stresses of higher magnitude [20]. While the number of cycles per day *v*_*n*_ has virtually no influence on bone volume fraction (Figure 4a), it is a strong inducer of bone damage (Figure 4b). This finding lends support to previous studies that have observed increased rates of musculoskeletal injury in horses that accumulate greater distances during training, at both canter and gallop speeds [6, 7].

The sensitivity of bone damage to both stress and cycles per day are subject to strong interactions between parameters (Figure S7b, electronic supplementary material). In particular, among pairs of parameters involving the cycles per day *v*_*n*_, the pair (*σ,v*_*n*_) has the highest second-order Sobol index (Figure S7c, electronic supplementary material). Since both high stress and high cycles per day are required for the formation of damage, it is conceivable that the effects on bone damage could be mitigated by restricting distances accrued at higher speeds. Thus, though we may speculate that ‘fast and light’ training programs (that are characterised by short program durations and lower cumulative gallop distances) [27] are less likely to result in injuries than higher volume programs, they were only associated with fewer injuries in two-year-old racehorses, and not mature racehorses [7]. A possible explanation for the lack of significant association in mature racehorses is the role of pre-existing injury and damage in causing future injuries, and that low grade injury is more tolerated in older racehorses.

The sensitivity values can have a range of interpretations, depending on the nature of each parameter. Parameters such as *v*_*n*_, *σ* and 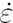 are external to the system and controllable through the design of training programs [57]. In this context, sensitivities characterise the effect of varying these parameters on our outputs of interest, namely the bone volume fraction *f*_*BM*_ and damage *D*^*∗*^. The other parameters are internal to the bone remodelling system and are often difficult or impossible to influence. Variations in these parameters may represent either uncertainties in the measurements of these parameters or biological heterogeneities inherent within the population. High sensitivities for these parameters point towards sources of uncertainty or variation in our estimates of model outputs [66]. We expect that gaining better confidence in these sensitive parameters (on either a population or individual level) through future studies would help us to model bone adaptation and damage more accurately. On the other hand, variation or uncertainty in insensitive parameters are less likely to be important in estimating *f*_*BM*_ and *D*^*∗*^, although we note that they may have consequential effects on other variables of interest. In our model, we find that the parameter *σ*_0_ has a large effect in simulating damage (Figure 4b). While this parameter is fitted to observational data by comparing to a median time to fracture, substantial variation exists between racehorses (Figure 3c,f). Thus, while microdamage is difficult to quantify in living racehorses (i.e. prior to catastrophic fracture), we believe that further work on understanding microdamage accumulation *in vivo* is warranted. We also find relatively high sensitivities in bone volume fraction to 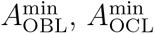 and *E*_0_ (Figure 4a). Thus, better estimates of bone formation and resorption rates would aid in the simulation of bone volume fraction. While bone stiffness *E*_0_ has been measured in racehorses, it greatly varies between horses [20], and incorporating the effects of this heterogeneity would be of interest for future work.

We use our model to simulate the response of bone volume fraction and damage in response to a dynamic training program including rest, progressive training and race-fit training (Figure 6). We find that fluctuations in bone volume fraction (Figure 6a) are consistent with those observed in horses resting and in training [24]. While bone microdamage is more difficult to measure, our model suggests that both progressive and race-fit training contribute to damage, with higher speed (*≥* 13.8 m/s) training accounting for the majority of damage (Figure 6f). Our model indicates that distances accumulated at canter speeds do not contribute substantially to damage (Figure 6f). However, prior studies have linked high canter distances to musculoskeletal injury when combined with high gallop distances [6]. This warrants further investigation, although we note that Verheyen et al. [6] categorise relatively fast speeds of up to 14 m/s as canter.

Our model simulations indicate that during training, bone damage does not plateau in time, and therefore all bones will eventually fail unless given the opportunity to repair damage through rest (Figure 6b). Holmes et al. [24] found lower rates of bone remodelling in training racehorses relative to resting racehorses, and on this basis recommend frequent rest to allow the bone to repair. While bone remodelling can occur during training, our model suggests that the repair associated with remodelling is insufficient to prevent the eventual failure of bone in response to prolonged training. In Victoria, Australia, rest periods can vary between 2–16 weeks, with the average reported to be 6.3 weeks in mature racehorses [7, 27]. Our model suggests that bone damage does not fully repair over a rest period of 6.3 weeks (Figure 6b), and therefore longer (or more frequent) rest periods are likely to benefit the repair of damage. We have simulated the response of bone to a typical training program from Victoria, Australia, where training volumes are believed to be higher than in the UK or US [27], although the durations and frequencies of rest tend to vary from region to region [67]. Rates of catastrophic musculoskeletal injury are higher in the UK and US compared to Australia [8], suggesting that workload alone likely does not explain variance in risk of injury between jurisdictions. The effect of other factors such as the duration and frequency of rest could be investigated in future work.

While previous models have coupled bone remodelling and damage [12, 13, 15, 42], we believe our study is the first to model these two processes dynamically in racehorses. Our model is calibrated to data on bone volume fractions and time to fracture from racehorses in training and rest. In contrast to the aforementioned studies [12, 15, 42], we do not observe reduced bone volume fraction in response to intense loading. This discrepancy is possibly because previous models account for the induction of bone remodelling in response to damage. We have not incorporated this process in our model (that is, Eq. (6) is independent of *D*^*∗*^) as third metacarpal bone volume fractions tend to increase near failure [68], possibly due to the inhibition of bone resorption at high loads [24]. However, focal bone resorption sites appear in response to damage, and are associated with fracture [48]. Further work is required to investigate the role of focal remodelling in bone injury. In the context of racehorses, Shaffer et al. [69] use a compartmental model to account for transitions between undamaged, damaged and resorbed bone. However, this model is analysed under the assumption of steady state and does not directly model the effects of exercise on transition rates. In contrast to our model, Shaffer et al. [69] find increased resorption rates associated with training in damaged regions of bone, possibly due to the appearance of focal remodelling at sites of fracture [48].

Given the cross-sectional nature of our data, there is substantial variation in the data used to fit our model (Figure 3), which we acknowledge as a limitation. The population of horses within the dataset exhibit variation in several characteristics known to be associated with catastrophic musculoskeletal injury [8] – such as age, sex, training practices and racing surfaces – that may also affect the dynamics of bone adaptation and microdamage [7, 70–72]. Although it would be valuable to examine how heterogeneities within the racehorse population might affect our findings, we do not believe the data used is sufficient to allow for such an analysis. Parameters such as osteoblast and osteoclast activity, as well as training speed and distance, are known to vary with age [27, 73]. For the purposes of this study, we have assumed these parameters to be constant in time (Figure 6). However, future work could incorporate age-varying training programs using training information reported by racehorse trainers [27].

As with our previous work [18], this study is limited by the use of a lumped parameter model. While we employ a lumped parameter model for computational efficiency, joint stresses, microdamage and repair processes are not evenly distributed throughout the bone [74–76]. Bergstrom et al. [77] have found associations between parasagittal groove fractures with more frequent rest and increased loads prior to bone injury. It would be of interest to incorporate our bone adaptation and damage equations into a finite-element model of bone mechanics to investigate these spatial heterogeneities [36, 39]. We also recognise that biochemical signalling and evolution of osteoblast and osteoclast densities may play a role in bone dynamics, causing delays in the response of bone to exercise [78, 79] and also providing the tissue with memory of the stresses incurred during the preceding days [80–82]. Accounting for these processes will enable better quantification of the response of bone to training programs that rapidly alternate between intense and less intense exercise. For simplicity, we have made the assumption that bone stiffness is independent of damage. While the stiffness of subchondral bone is well maintained for the majority of its fatigue life [20], this loss of stiffness could lead to feedback on the dynamics of bone remodelling. Finally, substantial variability exists on the level of both horses and trainers [27]. More detailed training and bone material property data is required to quantify heterogeneity in the bone adaptation parameters across populations of racehorses and to understand the effect of different training programs on bone injury.

## 5 Conclusion

We have developed a new mathematical model that couples the processes of bone adaptation and damage in the metacarpal subchondral bone of Thoroughbred racehorses. Our model is consistent with measurements of bone volume fraction and damage in response to rest and training. Simulations of the model indicate that joint stresses and the number of cycles per day are among the most important parameters in determining bone damage. Over the course of a typical racing preparation in Victoria, Australia, our model estimates that cycles accumulated at racing speeds are responsible for the majority of the damage. Our findings suggest that extended or more frequent rest periods are likely to be beneficial in repairing bone damage with the aim of preventing fractures in racehorses.

## 5.3 Data availability

The code associated with this manuscript is available from https://github.com/mic-pan/equine_bone_fatigue and is archived on Zenodo at https://doi.org/10.5281/zenodo.15745023 [83]. Since the data is owned by proprietary bodies and not available for distribution, the code for model calibration has been redacted.

