## Supplementary material for "A mathematical model of metacarpal subchondral bone adaptation, microdamage, and repair in racehorses"

Michael Pan, Fatemeh Malekipour, Peter Pivonka, Ashleigh V Morrice-West,
Jennifer A Flegg, R Chris Whitton, Peta L Hitchens

---

**C Supplementary figures** **11**

### A Supplementary methods

#### A.1 Derivation of bone repair equation

Since the damage $D^*$ ranges from 0 to 1, we treat this variable as the concentration of microcracks within the volume of bone mineral. Then, the total number of microcracks within the bone $X$ is defined as

$$X = f_{BM} D^*. \tag{S1}$$

The rate of microcrack formation and removal can then be treated as analogous to a chemical transport system, where the bone mineral is treated as a fluid, damage as the molecules of a chemical and flow rates as bone formation and resorption processes (Figure S1). From this figure, we see that

$$\frac{dX}{dt} = X'_D - X'_C, \tag{S2}$$

where $X'_D$ is the rate of microdamage accrual through loading (top of Figure S1) and $X'_C$ is the rate of microdamage removal through resorption (right of Figure S1). We assume that newly formed bone is free of damage, thus bone formation neither adds nor removes microcracks from the bone. From Eqs (8) and (S1),

$$X'_D = f_{BM} D'_f = f_{BM} v_D(\sigma) v_n(t). \tag{S3}$$

Under the assumption of the random appearance of remodelling processes, the rate of microcrack removal through resorption is equal to the product of microcrack concentration $D^*$ and resorption rate $A_{\text{OCL}}\alpha S_v$. However, in reality, osteoclasts tend to target microdamage, thus we multiply this expression by a repair specificity factor $F_s$. Thus,

$$X'_C = F_s A_{\text{OCL}} \alpha S_v D^* \tag{S4}$$

and

$$\frac{dX}{dt} = f_{BM} v_D(\sigma) v_n(t) - F_s A_{\text{OCL}} \alpha S_v D^*. \tag{S5}$$

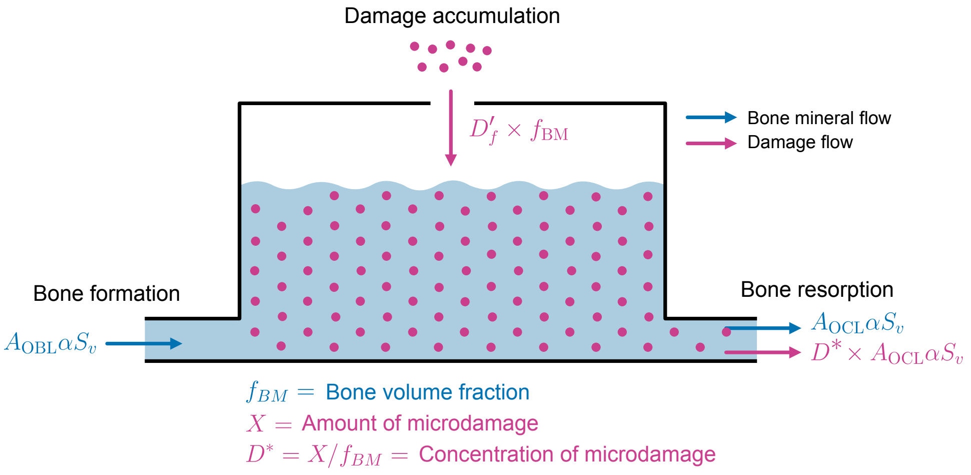

**Figure S1:** Schematic of damage formation and repair model, using a chemical transport analogy. Microcracks $X$ are indicated by dots, with their concentration given by $D^* = X/f_{BM}$. The fraction of bone mineral $f_{BM}$ is analogous to fluid. The deposition of bone (inflow) is assumed to be free of damage, whereas resorption removes microdamage at a rate proportional to the concentration of microcracks.

Applying the product rule to Eq. (S1),

$$\frac{dX}{dt} = \frac{dD^* f_{BM}}{dt} = \frac{dD^*}{dt} f_{BM} + D^* \frac{df_{BM}}{dt}. \tag{S6}$$

Hence, the net rate of damage accumulation is

$$\frac{dD^*}{dt} = \frac{1}{f_{BM}} \left( \frac{dX}{dt} - D^* \frac{df_{BM}}{dt} \right) \tag{S7}$$

$$= \frac{1}{f_{BM}} \left( f_{BM} v_D(\sigma) v_n(t) - F_s A_{\text{OCL}} \alpha S_v D^* - D^* A_{\text{OBL}} \alpha S_v + D^* A_{\text{OCL}} \alpha S_v \right) \tag{S8}$$

$$= v_D(\sigma) v_n(t) - \left[ A_{\text{OBL}} + (F_s - 1) A_{\text{OCL}} \right] \frac{\alpha S_v D^*}{f_{BM}}. \tag{S9}$$

By matching coefficients with Eq. (7),

$$D'_r = \left[ A_{\text{OBL}} + (F_s - 1) A_{\text{OCL}} \right] \frac{\alpha S_v D^*}{f_{BM}}, \tag{S10}$$

as per Eq. (12).

#### A.2 Training programs

##### *A.2.1 Influence of training speeds and distance on bone*

Our bone adaptation and damage model requires three inputs relating to racehorse training programs: the joint stress $\sigma$ (for Eqs (3),(11)), the strain rate $\dot{\varepsilon}$ (for Eq. (4)) and cycles (strides) per day $v_n$ (for Eq. (8)). Here, we show how these inputs relate to training variables, namely the speed $s$ [m/s] and distance $d$ [m] (Figure 2; green box).

We first express $\sigma$ and $\dot{\varepsilon}$ as continuous functions of the speed $s$. We relate the speed to the vertical ground reaction force $F_g$ [N/kg body weight] using an adapted quadratic equation from Witte et al. [1]:

$$F_g = 2.778 + 2.1376s - 0.0535s^2. \tag{S11}$$

Then, following Morrice-West et al. [2] and assuming that the maximum stress is $\sigma_{\text{max}} = 90$ MPa observed at the maximum speed of 20.99 m/s (corresponding to a maximum vertical force of $F_{\text{max}} = 24.13$ N/kg body weight), the vertical force is scaled to stress via

$$\sigma = \sigma_{\text{max}} \frac{F_g}{F_{\text{max}}}. \tag{S12}$$

Hitchens et al. [3] derived the strain rates for specific stress-states and their corresponding speeds and gaits. To extend this relationship to continuous speeds, we applied least squares fitting (using the speed and strain rate values in Table 3 of Hitchens et al. [3] as data points) to derive a quadratic equation to the strain rates for the stress-states $\sigma \leq 102$:

$$\dot{\varepsilon} = 0.01548s + 0.0004955s^2. \tag{S13}$$

Details of the quadratic fit can be found in Figure S2.

The stride length is calculated using the equations from Witte et al. [1]. In particular, the stride frequency $f$ [s$^{-1}$] is given by the quadratic equation

$$f = 1.7052 + 0.0305s + 0.0004s^2. \tag{S14}$$

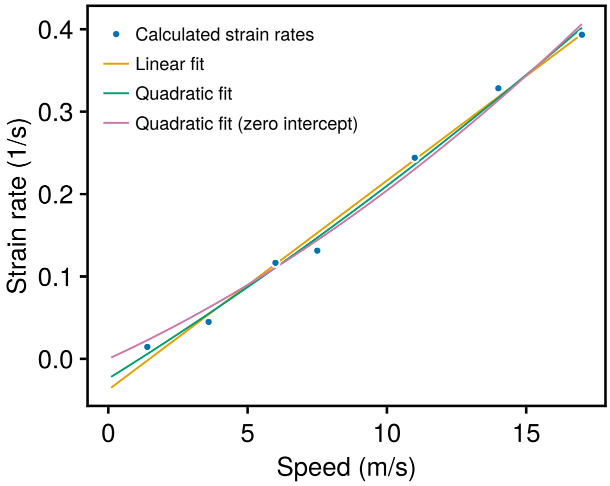

**Figure S2:** Fit of continuous functions to data on racehorse speeds and strain rates. The data (blue dots) are calculated using the equations in Table 3 of Hitchens et al. [3]. However, for consistency with our model, we use joint contact pressures calculated from Eqs (S11)–(S12), rather than the representative stresses from Table 3 of Hitchens et al. [3]. This results in strain rates of 0.01, 0.04, 0.12, 0.13, 0.24, 0.33 and 0.39 at the speeds 1.4, 3.6, 6.0, 7.5, 11, 14 and 17 m/s respectively. These data points are fitted using a linear function (yellow; $\dot{\varepsilon} = -0.03758 + 0.02539s$), quadratic (green; $\dot{\varepsilon} = -0.02396 + 0.02103s + 0.0002363s^2$) and quadratic with zero intercept (pink; $\dot{\varepsilon} = 0.01548s + 0.0004955s^2$). While both the linear and quadratic functions provide good approximations to the data, they predict negative strain rates at low speeds. We select the quadratic function with zero intercept for use in our model since it provides a comparable fit to the other functions while ensuring positive strain rates.

Therefore, the stride length $L$ [m] is estimated as

$$L = s/f. \tag{S15}$$

Given that a horse runs a distance of $d$ [m] per day at a particular speed, the corresponding cycles (strides) per day is

$$v_n = d/L. \tag{S16}$$

###### *A.2.2 Speeds and distances covered during training*

To simulate the response of bone to the range of training a three-year-old or older racehorse is subjected to during a preparation, defined as a period of training and/or racing between periods of rest, we define the speed and distance as functions of time (Figure 5, main text). In particular, we assume a typical training program in simulating the response of bone to training. We divide the preparation into four periods: rest (44 days, blue background), pre-training (28 days, blue background), progressive training (66 days, yellow background) and race-fit training (56 days, pink background) [4, 5]. Horses in rest (blue background) in an Australian training environment and in the absence of injury are typically turned out in a paddock with voluntary exercise including trotting for a distance of 4000 m every day [6], at a speed of 3.6 m/s [3]. The pre-training period (green background) consists of 2000 m/day at canter (7.5 m/s) [7]. Progressive training (yellow background) is divided into a slow phase (28 days, yellow box) and a fast phase (38 days, orange box) [4]. The slow phase consists of slower gallops of 11.8 m/s, linearly increasing to 13.8 m/s by the end of the four weeks. The galloping distance is also increased over this period, starting from 900 m/week before building up to 1800 m/week (the average galloping distance over the fast phase; Morrice-West et al. [4]) by the end of the slow phase. During the fast phase, racehorses cover a total distance of approximately 6600 m at 13.8 m/s and 3200 m at 16.0 m/s [4]. Over the 38 days of the fast phase, this results in an average gallop distance of 1805 m/week, which is assumed to be constant throughout the fast phase. Horses begin the fast phase galloping at 13.8 m/s, with the distance gradually replaced with the higher speed of 16.0 m/s. Assuming the distance per day at 16.0 m/s is linearly increased over this phase, horses cover 1179 m/week at 16.0 m/s by the end of progressive training. Horses in race-fit training (pink background) cover 4800 m at 13.8 m/s and 3200m at 16.0 m/s per month (red box), consistent with a medium-volume training program [4]. Weekly workloads are divided by seven to equate to an average daily workload. In addition, we model participation in an average race length of 1600 m every two weeks [4], at an average speed of 16.7 m/s [8] (dark red box). We include 4 races per preparation so that the racing preparation is 56 days and the total preparation length is 194 days, approximately consistent with the average rest frequency of 1.9 rests per year, or once every 192 days [4]. Jump outs and barrier trials (unofficial and official practice races typically 800m or less, respectively) are not considered since they typically replace race-fit training. Throughout both progressive and race-fit training, horses undergo constant 'slow workouts' amounting to 2000 m per day at a cantering speed of 7.5 m/s [3, 4].

Because racehorses train at several different speeds during a week, we combine their impacts on the strain energy densities sensed by cells and the rates of damage accumulation. More specifically, we assume that the osteoblast and osteoclast activities $A_{\text{OBL}}$ and $A_{\text{OCL}}$ respond to stresses associated with the highest speed at any given time point. Similarly, the strain rate $\dot{\varepsilon}$ is calculated at the highest speed (Eq. (S13)). However, the contribution of each speed is summed when calculating the rate of damage accumulation:

$$D'_f = \sum_{i=1}^{N} v_D(\sigma_i(t))v_{n,i}(t), \tag{S17}$$

where $\sigma_i$ and $v_{n,i}$ are the stresses and cycles per day corresponding to the $i$-th speed, and $N$ is the number of speeds considered.

#### A.3 Model calibration

##### A.3.1 Data

Two types of data are used to calibrate the mathematical model: (i) cross-sectional measurements of bone volume fraction in racehorses in training and rest, and (ii) the time (days) in training for horses that sustained fractures. Both data are sourced from previous experimental studies [9–11], which recorded information including bone volume fraction, time in training and cause of death (bone fracture or other causes). These datasets were subsequently combined in the modelling study by Hitchens et al. [3]. For (ii), we only consider mature horses ($\geq$ 3 years) that were in training when fracture occurred. The median time to fracture (133 days from start of progressive training) is used to fit the damage accumulation rate.

##### A.3.2 Objective function

Comparisons of the model to bone volume fraction and fracture under initial estimates for parameter values indicate discrepancies in the rates of bone formation and resorption, as well as resulting in a short modelled time to fracture. Given these discrepancies, we adjust the parameters $\mathbf{p} = (A_{\text{OBL}}^{\max}, A_{\text{OCL}}^{\max}, \sigma_0)$ to calibrate the model to these data. The remodelling rates $A_{\text{OBL}}^{\max}$ and $A_{\text{OCL}}^{\max}$ are used to adjust the bone formation and resorption rates. To adjust the rates of damage formation and resorption, we also fit the parameter $\sigma_0$ (representing a scaling factor for damage formation).

To assess the quality of fit to data, our model is simulated under three conditions:

1. **The adaptation of bone to training**. The model is simulated with the initial condition $f_{BM} = 0.878$ (the median volume fraction at rest), and a training speed of 16.0 m/s. Bone volume fractions are compared to the training data using the sum of squares function

$$J^{\text{train}} = \sum_{i=1}^{n_{\text{train}}} (f_{BM}^{\text{train}}(t_i^{\text{train}}) - f_{BM,i}^{\text{train}})^2, \tag{S18}$$

   where $(t_i^{\text{train}}, f_{BM,i}^{\text{train}})$ are the measurements ($n_{\text{train}} = 213$ data points) for horses in training from Hitchens et al. [3], and $f_{BM}^{\text{train}}(t)$ is the bone volume fraction recorded from the model.

2. **The de-adaptation of bone to rest.** The model is simulated with the initial condition $f_{BM} = 0.921$ (the median volume fraction in training) and a resting speed of 3.6 m/s. Bone volume fractions are compared to the resting data using the sum of squares function

$$J^{\text{rest}} = \left(\frac{n_{\text{train}}}{n_{\text{rest}}}\right)^2 \cdot \sum_{i=1}^{n_{\text{rest}}} (f_{BM}^{\text{rest}}(t_i^{\text{rest}}) - f_{BM,i}^{\text{rest}})^2, \tag{S19}$$

   where $(t_i^{\text{rest}}, f_{BM,i}^{\text{rest}})$ are the measurements ($n_{\text{rest}} = 24$ data points) for horses in rest from Hitchens et al. [3], and $f_{BM}^{\text{rest}}$ is the bone volume fraction recorded from the model. The scaling factor of $(n_{\text{train}}/n_{\text{rest}})^2$ is used to weight the training and resting datasets equally.

3. **The fracture of bone in response to high workload.** In simulating the time to fracture, we initialised bones to $f_{BM} = 0.878$ and simulated their response to progressive training followed by race-fit training. These programs are defined in Subsection A.2, however, we increase the monthly race-fit distances to 12,000 m at 13.8 m/s and 4,800 m at 16 m/s to simulate a high-volume workload [4]. Though this would not typically be warranted, to ensure the onset of fracture, the race-fit program is run for up to 365 days (rather than the 56 days specified in Figure 5). The time from the start of progressive training to fracture $t^{\text{frac}}$ is recorded, and compared to the data using the objective function

$$J^{\text{frac}} = (t_{\text{model}}^{\text{frac}} - t_{\text{median}}^{\text{frac}})^2, \tag{S20}$$

   where the median time to fracture is $t_{\text{median}}^{\text{frac}} = 133$ days.

We note that since conditions 1 and 2 only require the simulation of bone volume fractions, distances have not been defined for these training programs as the quality of fit is independent of training volume.

The full objective function used to calibrate the model is given by the weighted sum of squares function

$$J(\mathbf{p}) = J^{\text{train}} + J^{\text{rest}} + J^{\text{frac}}. \tag{S21}$$

##### A.3.3 Parameter optimisation

The optimal set of parameters is found using an adaptive differential evolution optimiser with radius limited sampling as a global optimiser [12]. The solution of the global optimiser is used as the starting point for a local optimiser (the Broyden-Fletcher-Goldfarb-Shanno algorithm; Nocedal and Wright [13]) to find the optimal value of $\mathbf{p}$.

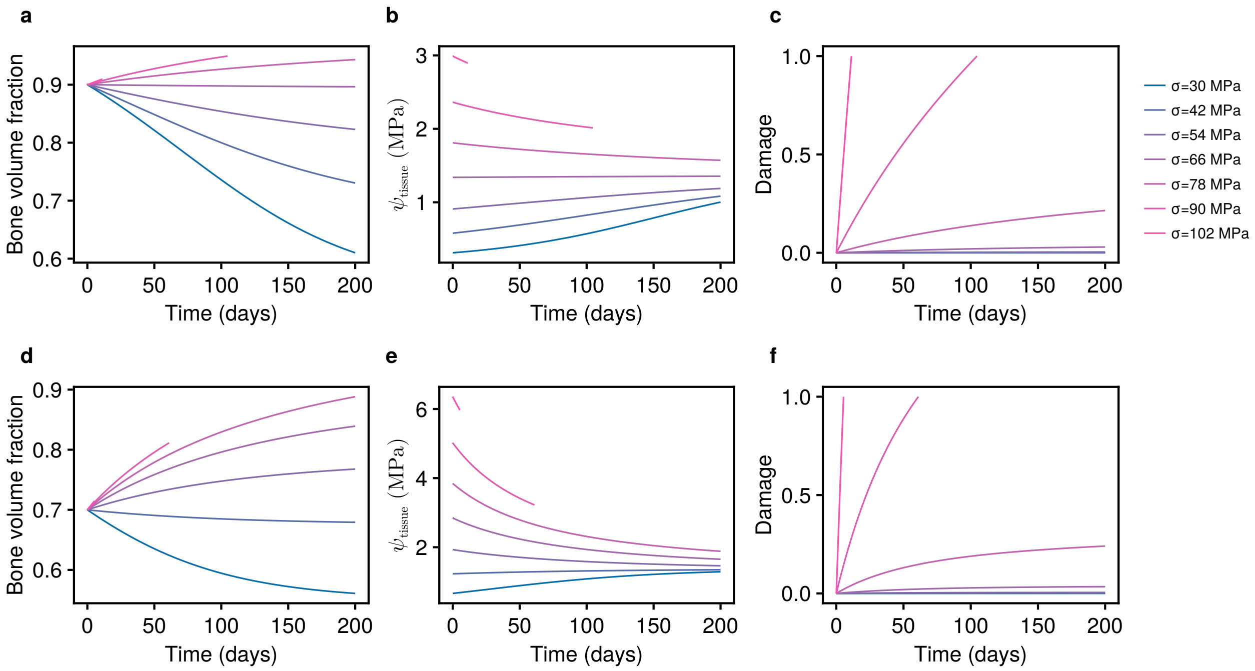

**Figure S3:** Simulations of bone adaptation and damage under varying stresses $\sigma$, run for $t = 200$ days. (a–c) Simulations using an initial condition with bone volume fraction $f_{BM} = 0.9$; (d–f) simulations using an initial condition of $f_{BM} = 0.7$. Stresses $\sigma$ are varied from 30 MPa to 102 MPa, and the corresponding strain rates from Table 3 of Hitchens et al. [3] are used. We investigate the effects of varying stress, independent of the cycles per day $v_n$. Accordingly, the cycles per day is set to a constant value of $v_n = 40$ day$^{-1}$, which corresponds to a distance of 240 m/day (1680 m/week) assuming a galloping stride length of approximately 6 m. Simulations are terminated if the damage variable $D^*$ reaches a value of 1, in which case the bone is considered to have failed.

##### A.4 Sensitivity analysis

To detect relationships between parameters and outputs of interest, a sensitivity analysis of the model is conducted by sampling parameters from the ranges shown in Table 1. Simulations are run for $t = 70$ days to emulate the average time from the start to end of a training program. The bone volume fraction $f_{BM}$ and damage $D^*$ are recorded at the end of the simulation. Sensitivity analyses are conducted using two methods: (i) the partial rank correlation coefficient (PRCC) [14] to detect the strength of associations between parameters and model outputs as well as the direction of association; and (ii) Sobol indices [15] that attribute the variance in model outputs to variances in parameters and their combinations.

To reduce the risk of detecting spurious relationships, we include a dummy variable in the sensitivity analyses that has no impact on the model output [14]. Samples are generated using Sobol sequences [16]. Parameter sensitivities are calculated using $N = 10^5$ samples for PRCC and $N = 10^5$ samples for each design matrix for Sobol indices.

#### B Supplementary results

##### B.1 Response of bone to different training speeds

Simulations of the calibrated mathematical model under different stresses are shown in Figure S3. As the stress $\sigma$ increases from 30 MPa to 90 MPa, the bone volume fraction at the end of

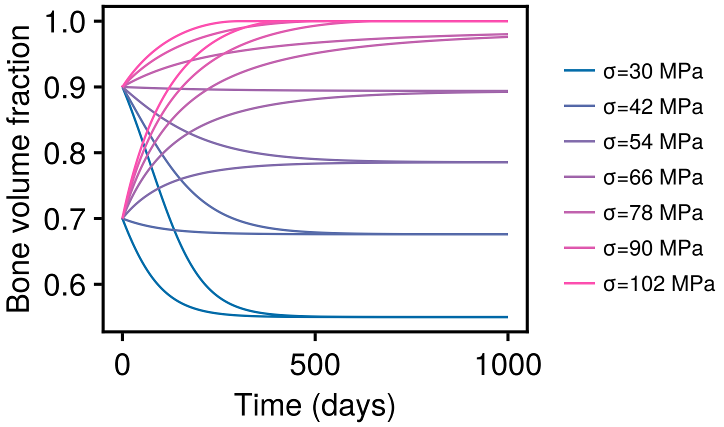

**Figure S4:** Simulations of bone adaptation to steady state. Simulations are run from two initial conditions, $f_{BM}(0) = 0.9$ and $f_{BM}(0) = 0.7$ under the same conditions as in Figure S3, but with simulations run to $t = 1000$ days. Lines are coloured by the applied stress.

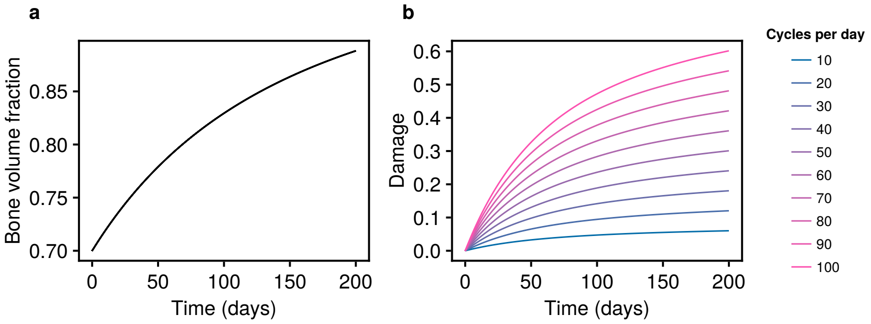

**Figure S5:** Simulations of bone volume fraction ($f_{BM}$) and damage ($D^*$) at different cycles (strides) per day ($v_n$). (a) Bone volume fraction (Eq. (2)). Only one line is displayed since the bone volume fraction evolves identically for all cycles accumulated per day. (b) Damage incurred by training (Eq. (7)), where $D^* = 1$ indicates bone failure. All simulations are run with a stress of $\sigma = 78$ MPa (corresponding to a slow gallop) and an initial bone volume fraction of $f_{BM} = 0.7$.

the simulation period is increased (Figure S3a), conferring better resistance to microdamage. However, higher stresses also induce greater damage, with the model predicting failure at the two highest stresses $\sigma = 90,\ 102$ MPa (Figure S3c).

We also compare the response of bone from an adapted state of $f_{BM} = 0.9$ (Figure S3a–c) to bone from an unadapted state of $f_{BM} = 0.7$ (Figure S3d–f). While the unadapted bone achieves a lower bone volume fraction by the end of the simulation when exposed to the same stresses, the steady-state bone volume fraction is the same regardless of initial condition, as illustrated by simulations of the same model over extended time periods (Figure S4). Our model shows that when bone is exposed to high stresses prior to adaptation, damage accumulates at faster rates. For instance, at the racing stress of $\sigma = 90$ MPa, the bone is predicted to fail at $t = 61$ days (Figure S3f), substantially less than the time taken for the bone to fail from an adapted state ($t = 105$ days; Figure S3c). While bone does not fail at the galloping stress of $\sigma = 78$ MPa over the simulation period, there is a clear period of accelerated damage accumulation in the first 50 days, before the bone volume fraction has adapted (Figure S3f).

Figure S5 shows the effect of number of cycles (strides) per day on damage accumulation. Since we do not model the effect of damage on bone stiffness, the bone volume fraction evolves

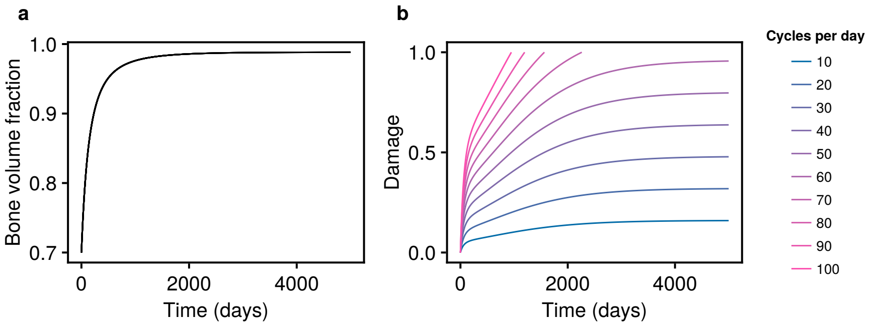

**Figure S6:** Long-term simulations of bone adaptation and damage at different cycles (strides) per day. (a) Bone volume fraction. Only one line is displayed since the bone volume fraction evolves identically for all cycles per day. (b) Damage incurred by training. All simulations are run with a stress of $\sigma = 78$ MPa (corresponding to a slow gallop) and an initial bone volume fraction of $f_{BM} = 0.7$.

identically for all cycling rates. However, the rate of damage accumulation increases in proportion to the number of cycles per day. While none of the cycles per day lead to failure over the durations simulated, the extent of bone remodelling is reduced at higher bone volume fractions due to the corresponding decrease in specific surface area $S_v$ (Eq. (1)). As a result, should the loading continue indefinitely, accumulating 70 or more cycles per day will eventually result in failure (Figure S6).

#### B.2 Sobol sensitivities

To quantify the magnitude of the effects on model outputs, we conduct a sensitivity analysis using Sobol indices (Figure S7). High Sobol indices indicate parameters with a strong influence on the output variables of interest ($f_{BM}$ and $D^*$). The parameters $\sigma$, $A_{\mathrm{OBL}}^{\mathrm{min}}$, $E_0$ and $A_{\mathrm{OCL}}^{\mathrm{min}}$ have the greatest effect on the bone volume fraction, consistent with the results from PRCC (Figure S7a). Similarly, $\sigma$, $\sigma_0$, $v_n$ and $E_{\mathrm{nom}}$ are the parameters with the greatest effect on bone damage (Figure S7b). However, $\sigma_1$, $E_0$ and $F_s$ have relatively modest total Sobol indices, in contrast to their high PRCC values.

While the parameter sensitivities on bone volume fraction are predominantly first-order, there are substantial higher order effects on damage (Figure S7b). This indicates that some combinations of parameters are important in explaining variations in damage, and we therefore display the second-order Sobol indices in Figure S7c, with damage on the top right. The most significant pair of parameters contributing to damage is $(\sigma, \sigma_0)$, due to their difference appearing in the exponential relationship in Eq. (10). The pair $(\sigma, v_n)$ (indicated by the red asterisk) is associated with a modest (but non-zero) second-order sensitivitity, indicating that both high speeds and (moderately) high distances lead to increased bone damage, beyond the sum of their individual effects.

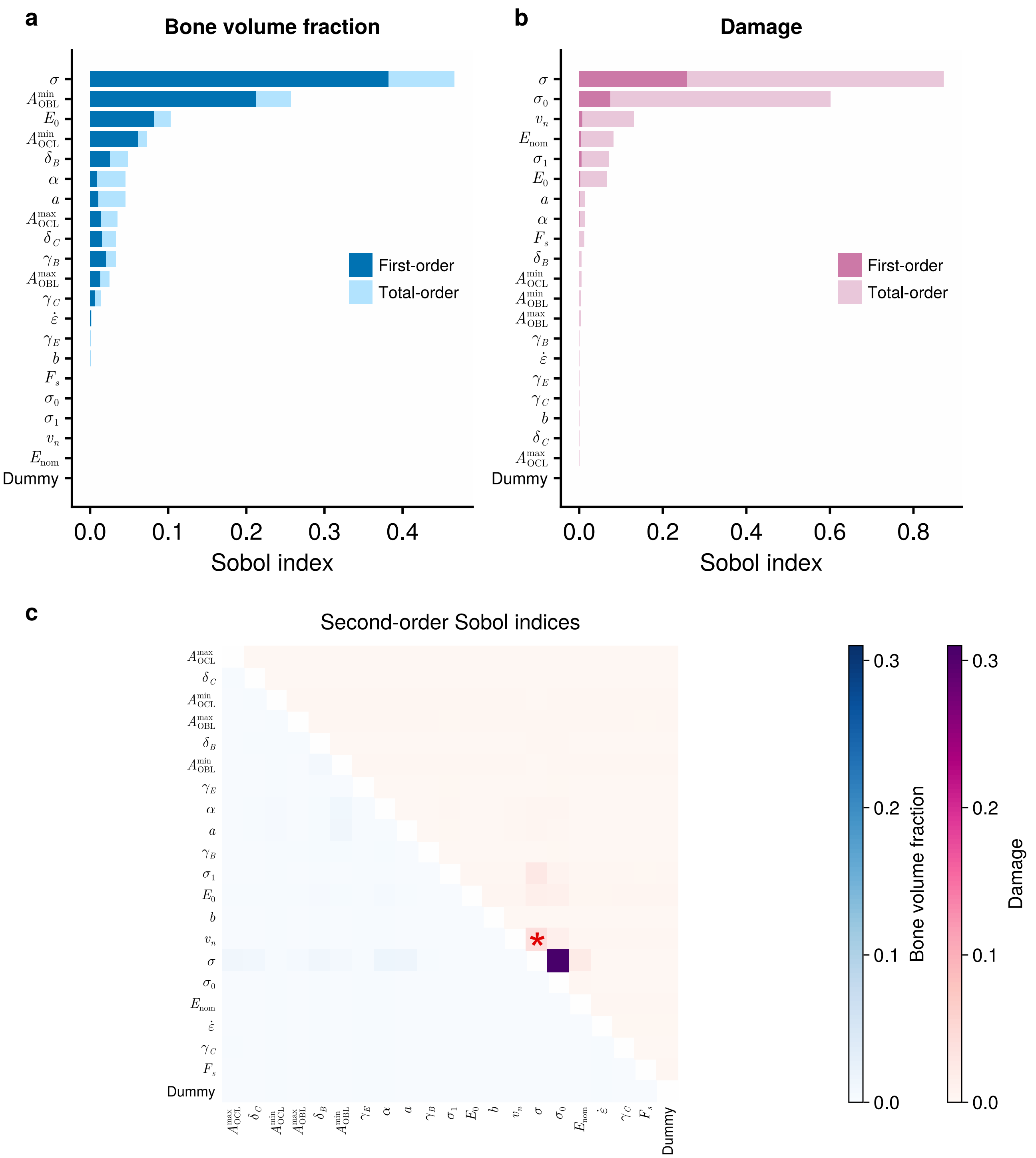

**Figure S7:** Sensitivity analysis of parameters using Sobol indices. Effects of each parameter on (a) bone volume fraction $f_{BM}$; and (b) damage $D^*$ after 10 weeks. High Sobol indices indicate parameters with a strong influence on the output variables of interest. The model described by Eqs (1)–(12) is simulated using an initial condition of $f_{BM} = 0.7$ and $D^* = 0$. Descriptions of parameters and their ranges are shown in Table 1. (c) Second-order Sobol indices for both bone volume fraction (lower left) and damage (upper right). High Sobol indices indicate parameters with a strong influence on the output variables of interest. The index corresponding to the effect of pair $(\sigma, v_n)$ on damage is indicated by the red asterisk.

#### C Supplementary figures

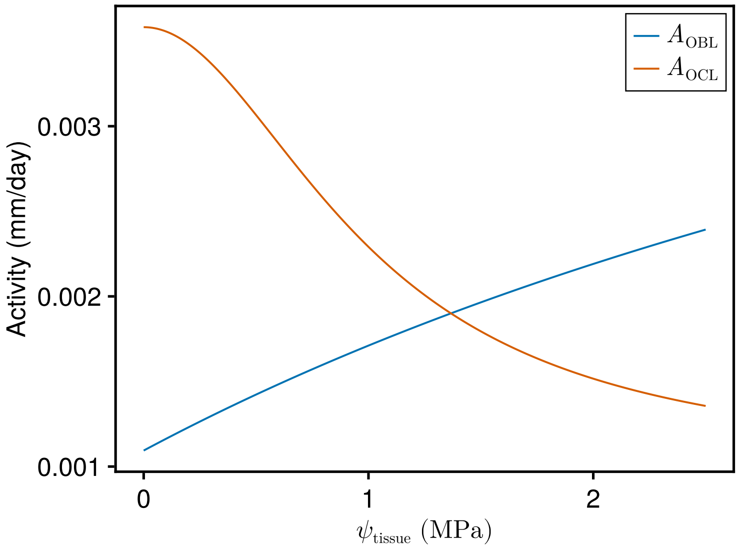

**Figure S8:** Relationship between strain energy density $\psi_{\text{tissue}}$ and osteoblast ($A_{\text{OBL}}$) and osteoclast ($A_{\text{OCL}}$) activities. $A_{\text{OBL}}$ and $A_{\text{OCL}}$ are calculated using Eqs (5),(6), using parameters obtained from fitting to measurements of bone volume fraction in racehorses (Table 1). The intersection of the curves at $\psi_{\text{tissue}} = 1.37$ MPa represents the setpoint, which is consistent with loading states associated with trotting and cantering (3.6–7.5 m/s; Table 3 of Hitchens et al. [3]).

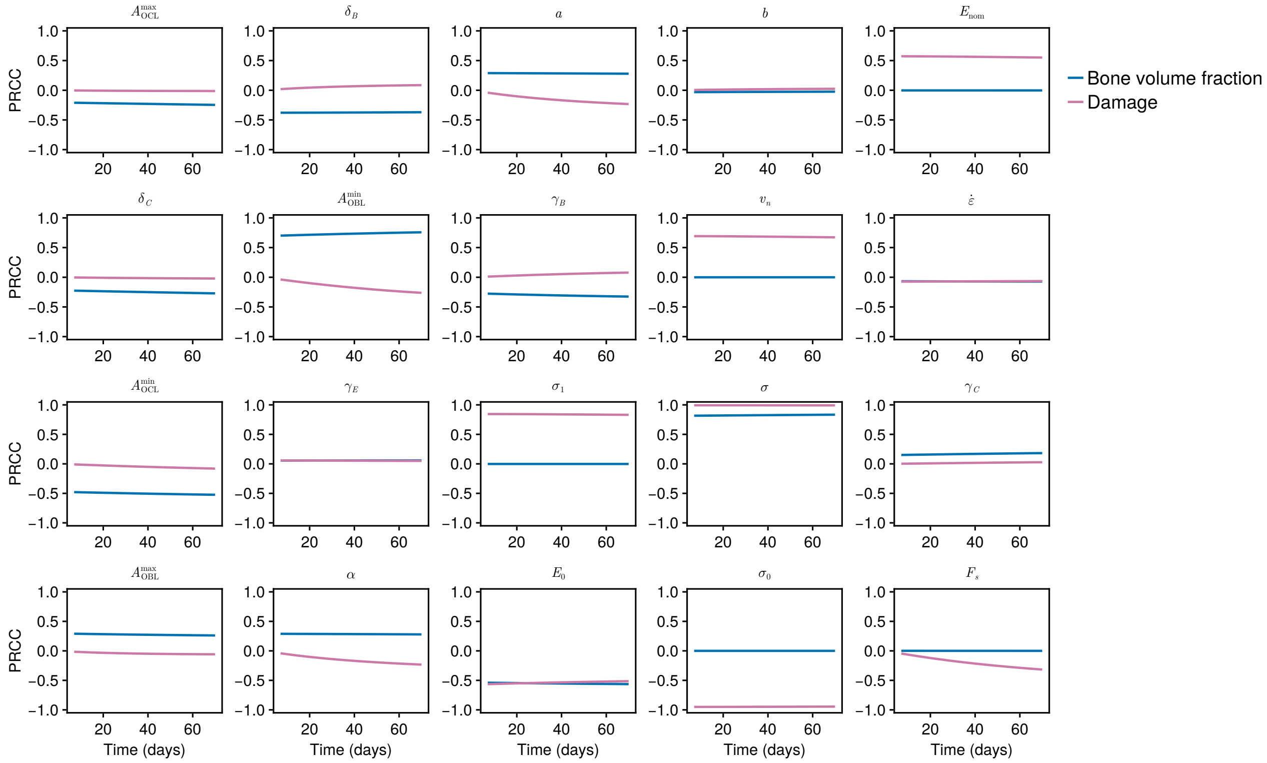

**Figure S9:** PRCC values for each parameter at different points in time. Simulations of the model are run under constant inputs for 70 days, and sensitivities are calculated at 7-day intervals.

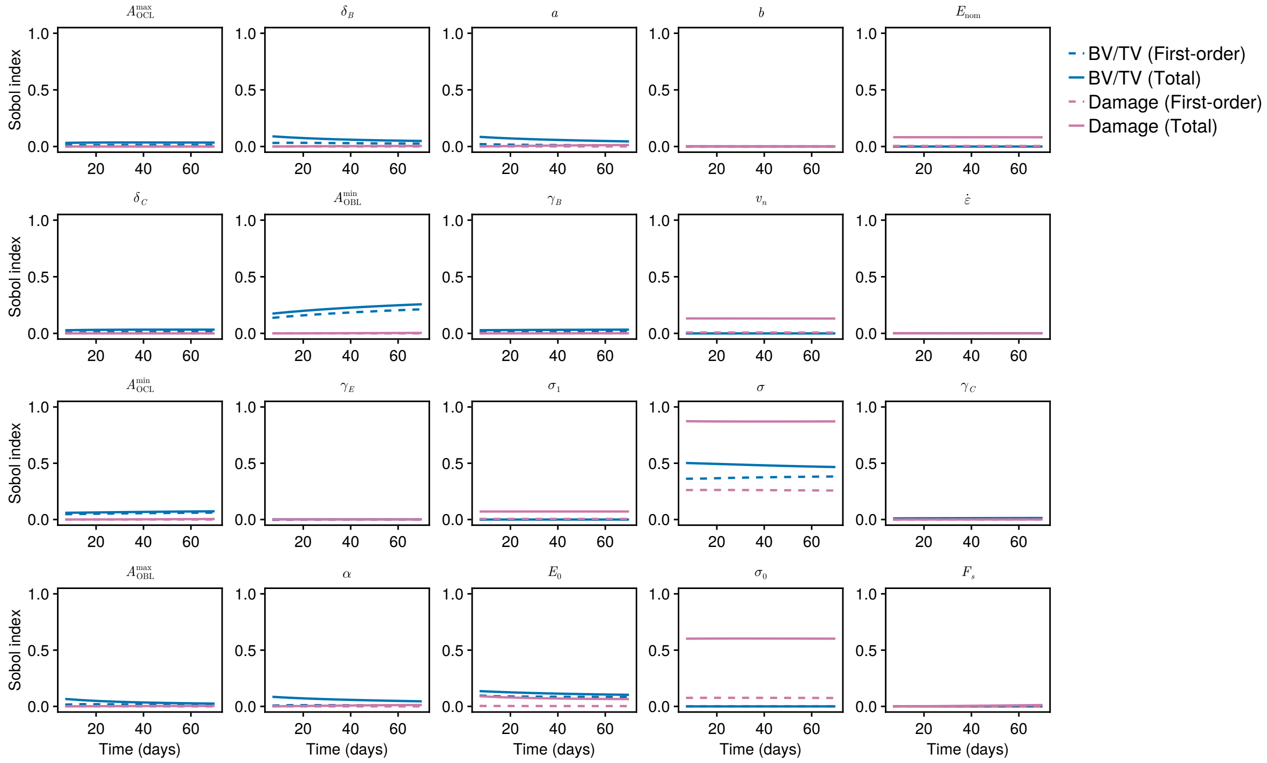

**Figure S10:** First-order and total order Sobol sensitivities for each parameter at different points in time. Simulations of the model under constant inputs are run for 70 days, and sensitivities are calculated at 7-day intervals.
